# Effect of *Ascosphaera apis* infestation on the activities of four antioxidant enzymes in Asian honey bee larval guts

**DOI:** 10.1101/2022.12.27.522003

**Authors:** Kaiyao Zhang, Zhongmin Fu, Xiaoxue Fan, Zixin Wang, Siyi Wang, Sijia Guo, Xuze Gao, Haodong Zhao, Xin Jing, Peiyuan Zou, Qiming Li, Mengjun Chen, Dafu Chen, Rui Guo

## Abstract

*Ascosphaera apis* exclusively infects bee larvae and causes chalkbrood, a lethal fungal disease that results in the sharp reduction in adult bees and colony productivity. However, little is known about the effect of *A. apis* infestation on the activities of antioxidant enzymes in bee larvae. Here, *A. apis* spores were purified and used to inoculate Asian honey bee (*Apis cerana*) larvae, followed by detection of the host survival rate and evaluation of the activities of four major antioxidant enzymes. At 6 days post inoculation (dpi) with *A. apis* spores, white mycelia penetrated the posterior end of the larva, extended to the anterior end, and eventually covered the entire larval body surface, presenting an obvious symptom of chalkbrood disease similar to that occurs in *Apis mellifera* larvae. Additionally, PCR identification showed that the expected fragment was amplified from the *A. apis*-inoculated larval guts and the *A. apis* spores, verifying the *A. apis* infection of *A. cerana* larvae. The survival rate of larvae inoculated with *A. apis* was high at 1–2 dpi, sharply decreased to 4.16% at 4 dpi, and reached 0% at 5 dpi; whereas that of un-inoculated larvae was always high at 1~8 dpi, with an average survival rate of 95.37%, indicating the negative impact of *A. apis* infection on larval survival. Furthermore, in comparison with those in the corresponding un-inoculated groups, the superoxide dismutase (SOD) and catalase (CAT) activities in the 4-day-old larval gut in the *A. apis*-inoculated groups were reduced (*p* > 0.05), while those in the 5- and 6-day-old larval guts were significantly decreased (*p* < 0.05); the glutathione S-transferase (GST) activity in the 4- and 5-day-old larval guts was significantly increased (*p* < 0.05), while that in the 6-day-old larval gut was reduced (*p* > 0.05); the polyphenol oxidase (PPO) activity in 4-day-old larval gut was increased (*p* > 0.05) and that in the 5-day-old larval gut was significantly increased (*p* < 0.05), whereas that in the 6-day-old larval gut was significantly reduced (*p* < 0.01). These results together suggested that the activities of SOD and CAT in the larval guts were suppressed during the process of *A. apis* infestation, while the GST activity was induced to activation, and the PPO activity was first enhanced and then inhibited. Our findings not only unravel the response of *A. cerana* larvae to *A. apis* infestation from a biochemical perspective, but also offer a valuable insight into the interaction between Asian honey bee larvae and *A. apis*.

## 1. Introduction

Honey bees are of great importance due to their pollination of numerous wildflowers and agricultural crops, production of *api*-products such as royal jelly and propolis, and scientific model applications [1]. However, as a kind of representative eusocial insect, honey bees are prone to infection by pathogens and parasites [2,3]. Among these, *Ascosphaera apis*, an obligate fungal pathogen of honey bee brood, causes chalkbrood disease, which results in a dramatic decline of colony strength and productivity alone or in combination with other biotic or abiotic factors [4]. For insects, including honey bee, the midgut is not only a pivotal tissue responsible for food digestion, enzyme secretion, and nutrient absorption, but also a primary site of interaction with pathogenic microorganisms [5].

The fungal pathogen invades the body wall and then multiplies in the blood cavity, leading to the death of the host. After long-term interaction and coevolution, insects have evolved various mechanisms to resist pathogenic infestation, including physical and chemical barriers and an innate immune system [6]. Insects such as honey bee can produce reactive oxygen species (ROS)—a by-product of aerobic metabolism—to defend themselves against xenobiotics or pathogens, thus maintaining the stability of their internal environment [6]. All aerobic organisms, including insects, have specific antioxidant systems to scavenge excess oxygen radicals in the body to avoid oxidative damage [7]. Studies have shown that organisms protect themselves from oxidative damage by regulating intracellular ROS production through a system of antioxidant enzymes, including superoxide dismutase (SOD), catalase (CAT), and glutathione S-transferase (GST) [8–14]. Recently, Li et al. [10] found that *Ascosphaera apis* infestation induced oxidative stress in *Apis mellifera* worker larvae, resulting in a significant decrease in the enzymatic activities of SOD, CAT, and GST.

*Apis cerana*, a bee species distributed throughout various climatic zones of Asia, is predominant and widely used in the Asian beekeeping industry and plays a critical role in local pollination and ecological maintenance [15,16]. In beekeeping, *A. mellifera* larvae are commonly liable to *A. apis* infection, and *A. cerana* larvae often die from chalkbrood. Previously, our group isolated *A. apis* from chalkbrood mummies of *A. cerana* and demonstrated that inoculation of *A. cerana* larvae with *A. apis* spores under lab conditions resulted in chalkbrood disease [17]. However, whether *A. apis* invasion affects the activities of major antioxidant enzymes is still largely unknown.

In the current work, *A. apis* spores were purified and used to inoculate the *A. cerana* 3-day-old larvae, followed by calculation of the host survival rate and the activities of four antioxidant enzymes such as SOD, CAT, GST, and polyphenol oxidase (PPO) in the guts of both *A. apis*-inoculated and un-inoculated larvae. Our data clarify the effect of *A. apis* invasion on the activities of the aforementioned antioxidant enzymes and offer a valuable understanding of *A. apis* response of *A. cerana* larvae.

## 2. Materials and Methods

### 2.1. Purification of *A. apis* spores

*A. apis* stored at 4 °C was transferred to PDA medium and cultured at 33 ± 0.5 °C in a constant temperature and humidity chamber (Jingke, China). Ten days after culturing, white mycelia were removed, and black fruiting bodies were harvested and transferred to an RNAase-free EP tube following our previously established protocol [18,19]. Next, 1 mL of sterile water was added to the EP tube, followed by complete grinding. Then, the grinding fluid was centrifuged at 25 °C, 7000 × g for 3 min. The supernatant was removed, and 1 mL of sterile water was added, followed by centrifugation at 25 °C, 7000 × g for 3 min. The centrifugation was repeated twice to clean the spores, which were then stored at 4 °C until use.

### 2.2. Experimental inoculation and survival rate calculation

Honey bee larvae were reared and inoculated with *A. apis* spores based on our previously described method [20]. In brief, (1) the larvae diet was prepared following Feng et al. [21], preheated to 35 °C, and added to 6-well culture plates. (2) PCR amplification was performed to detect the honey bee colonies reared in the apiary, and three colonies with negative results were selected as experimental colonies. The 2-day-old larvae were carefully transferred to 6-well culture plates using a Chinese graft, and after 24 h, the larvae were transferred to 48-well culture plates (1 larva/well) and placed in an incubator (35 °C, 95% RH). (3) The purified *A. apis* spores were subjected to gradient dilution and mixed with the diet, with a final concentration of 1×10^7^ spore/mL. The 3-day-old treatment group larvae were fed the diet containing spores, while 3-day-old larvae in the control groups were fed the diet without spores; the diet was changed daily. There were three biological replicas of this experiment. The dead larvae in both treatment and control groups were recorded and removed every 24 h until 8 dpi, followed by survival rate calculations.

The 4-, 5-, and 6-day-old larvae guts (inoculated with A. apis spores and un-inoculated) were dissected, frozen in liquid nitrogen, and held at −80 °C until use.

### 2.3. Evaluation of the SOD activity

The larval gut SOD activity was evaluated using the Insect SOD ELISA Kit (MLBIO, China). Briefly, the gut samples were fully ground with a high-throughput tissue grinder (MEIBI, China); the grinding fluid was transferred to a sterile EP tube; 750 mL of 1×PBS solution was added, followed by centrifugation at 1000 × g for 10 min, the supernatant was incubated on ice; 50 mL of the standard sample was added to the standard well, while 40 mL of sample diluent and 10 mL of grinding fluid were added into the sample well followed by gentle shaking. Meanwhile, a blank well was set, sealed with a film, and the microtiter plate was incubated at 37 °C for 30 min. The reaction solution was then discarded, and washing buffer was added to the wells. The solution was removed after standing for 30 s, and the operation was repeated 5 times. Then, 50 mL of enzyme standard reagent was added to each standard well and sample well, and 50 mL each of chromogenic A and B were added. After gentle shaking, the solution was placed at 37 °C for 10 min; 50 mL of termination solution was added into each well to terminate the reaction; finally, the OD value at 450 nm from each well was measured by a Thermo Scientific™ Varioskan™ LUX (ThermoFisher, USA). This experiment included three biological replicas.

The specific antioxidant enzyme activity was expressed as units of enzyme activity per milligram of protein. Data were analyzed and plotted by GraphPad Prism 8 software (GraphPad Software, San Diego, California). Experimental data were presented as Mean ± SD and subjected to Student’s *t*-tests.

### 2.4. Examination of CAT and GST activities

The larval gut CAT activity was examined with an Insect CAT ELISA Kit (MLBIO, China); GST activity was with an Insect GST ELISA Kit (MLBIO, China). The operation and calculation methods were the same as in section 2.3.

### 2.5. Detection of PPO activity

The gut samples (each 0.1 g) in the six groups mentioned above were transferred to sterile EP tubes, and 1 mL of extraction solution was added. Next, the gut tissues were ground thoroughly using a high-throughput tissue grinder, followed by centrifugation at 8000 × g for 10 min. The supernatant was transferred to a new EP tube and placed on ice for measurement. The assay tube and control tube reaction systems were prepared, placed in a 25 °C water bath for 10 min, and then quickly transferred to a 100 °C metal bath for 10 min. The reaction system was mixed thoroughly and centrifuged at 5000 × g for 10 min, and the supernatant was transferred to a new EP tube and placed on ice. The assay and control OD values were respectively named A assay and A control, and the difference between them was named ΔA. PPO activity was calculated as: PPO (U/g) = 120×ΔA÷W (ΔA = A assay − A control), W: sample mass, g.

## 3. Results

### 3.1. Verification of *A. cerana* larvae infection by inoculation with *A. apis* spores

Here, a prominent symptom of chalkbrood disease was observed in larvae inoculated with *A. apis* spores—white mycelia first penetrated from the posterior end of the larva at 6 dpi, extended to the anterior end, and eventually covered the entire larval body surface (Figure 1A). Additionally, as shown in Figure 1B, agarose gel electrophoresis indicated that fragments with the expected sizes (about 217 bp) could be amplified from the *A. apis*-inoculated larval guts and *A. apis* spores, but could not be amplified from the un-inoculated larval guts and sterile water. Collectively, these results confirmed the infection of *A. cerana* larvae by inoculation with *A. apis* spores.

**Figure 1.**
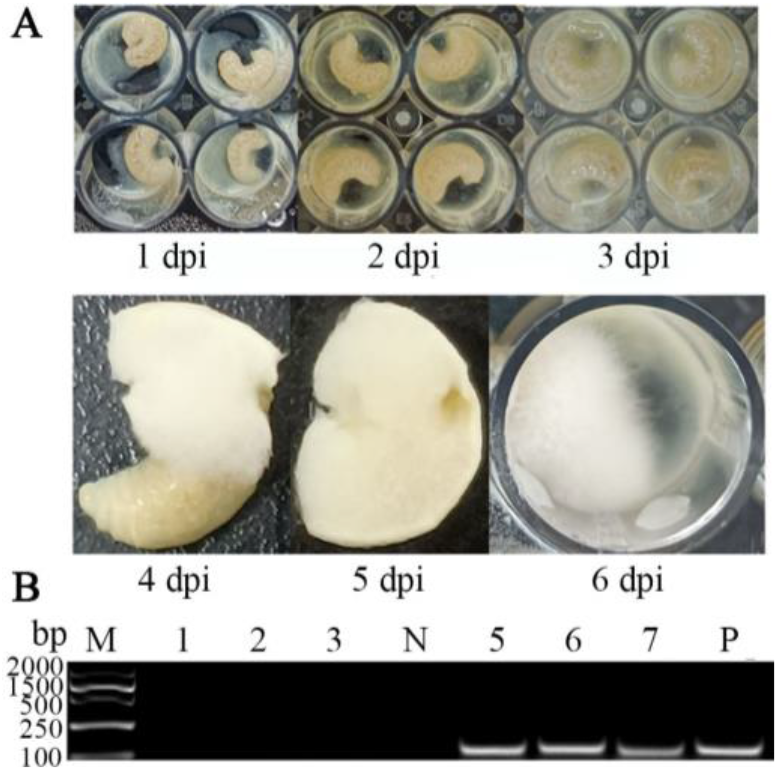
Observation and verification of *A. apis* infection of *A. cerana* larvae. (**A**) Observation of larval chalkbrood symptom after *A. apis* spore inoculation; (B) PCR validation of larval guts inoculated with *A. apis* spores. Lane M: DNA marker; Lane 1–3: Un-inoculated 6-, 5-, and 4-day-old larval guts; Lane N: Sterile water (negative control); Lane 5–7: *A. apis*-inoculated 6-, 5-, and 4-day-old larval guts; Lane P: Purified spores of *A. apis* (positive control).

### 3.2. Survival rate of *A. c. cerana* larvae after *A. apis* spores infection

All figures and tables should be cited in the main text as Figure 1, Table 1, etc. The survival rate of *A. cerana* larvae in the *A. apis*-inoculated group were 97.92%, 83.33%, and 33.33% at 1 dpi~6 dpi, respectively; the larval survival rate sharply decreased to 4.16% at 4 dpi, and 0 at 5 dpi. Comparatively, the survival rate of larvae in the un-inoculated group was always high at 1 dpi~8 dpi, with an average survival rate of 95.37% (Figure 2).

**Figure 2.**
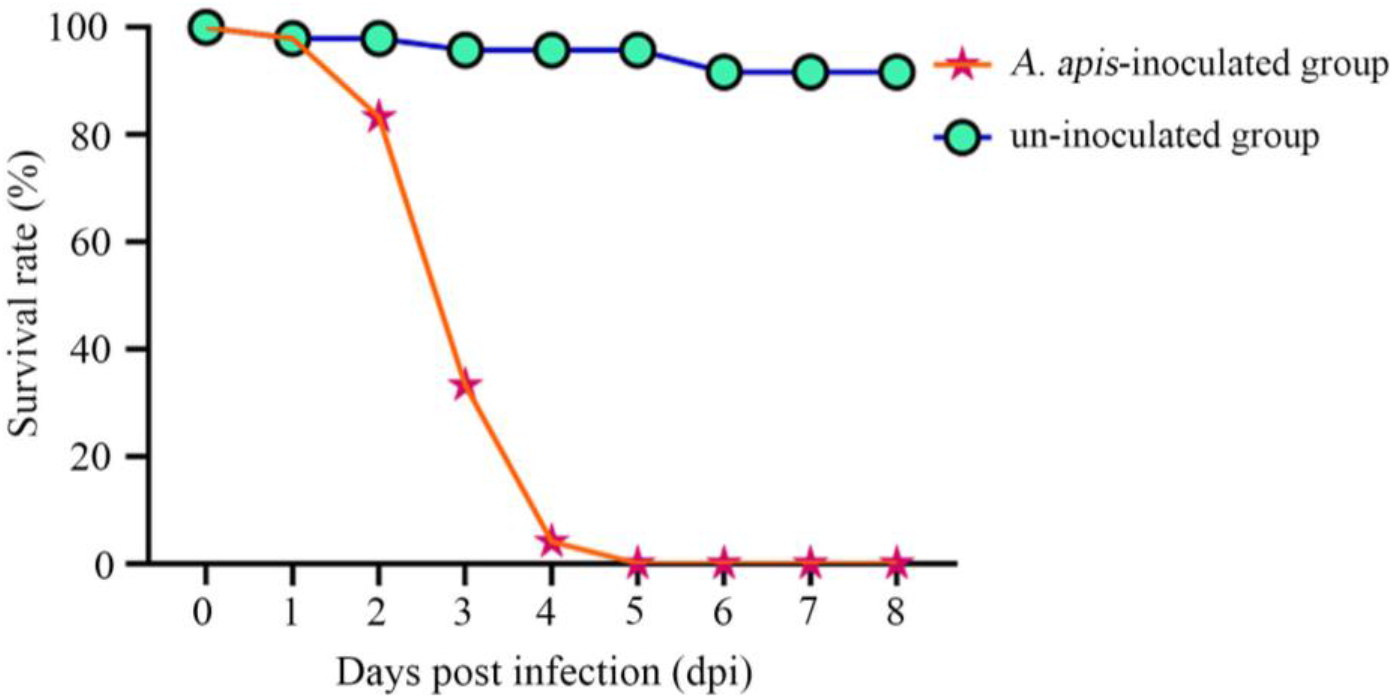
Survival rates of *A. apis*-infected and un-infected *A. cerana* larvae.

### 3.3. Effect of A. apis infection on SOD activity in A. cerana larval guts

As compared with the corresponding un-infected groups, the SOD activity in the 4-day-old (1.93 U/mL) larval gut in the *A. apis*-infected group was reduced (*p* > 0.05), while that in the 5(2.86 U/mL) and 6-day-old (2.79 U/mL) larval guts was significantly decreased (*p* < 0.05) (Figure 3).

**Figure 3.**
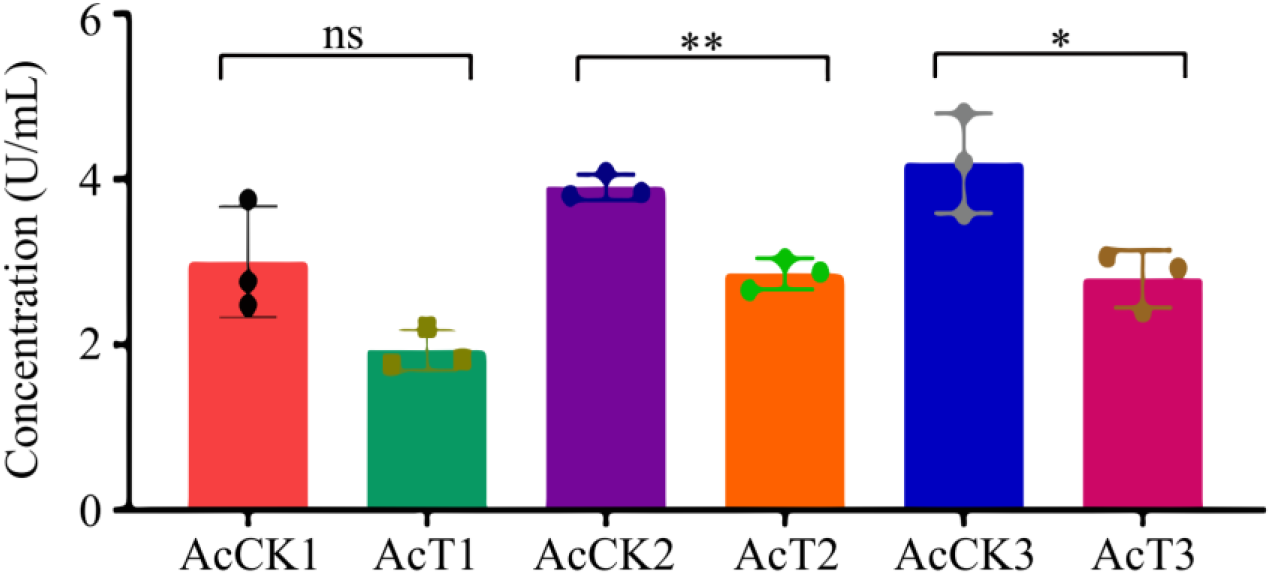
SOD activity in *A. cerana* 4-, 5-, and 6-day-old larval guts infected by *A. apis*. The experimental data are shown as Mean ± SD and subjected to Student’s *t*-tests, ns: *p* > 0.05, *: *p* < 0.05, **: *p* < 0.01.

### 3.4. Effect of *A. apis* infection on the CAT activity in the *A. cerana* larval guts

In comparison with the corresponding un-infected groups, CAT activity in the 4-day-old (1.63 U/mL) *A. apis*-infected larval gut was decreased (*p* > 0.05), whereas that in the 5- (1.54 U/mL) and 6-day-old (2.91 U/mL) larval guts was significantly reduced (*p* < 0.05) (shown in Figure 4).

**Figure 4.**
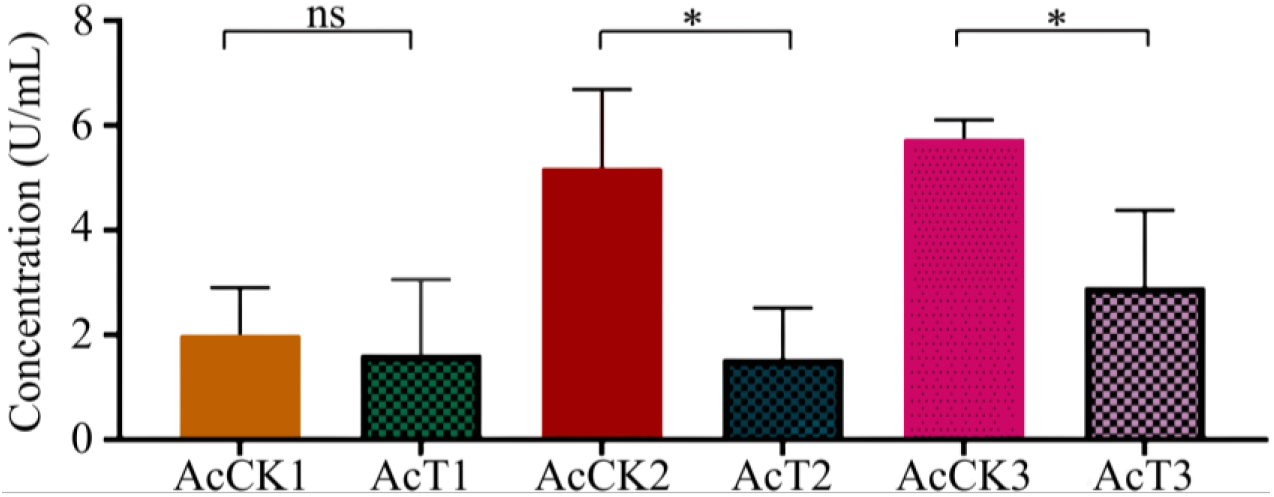
CAT activity in *A. cerana* 4-, 5-, and 6-day-old larval guts infected by *A. apis*.

### 3.5. Effect of *A. apis* infection on GST activity in *A. cerana* larval guts

Compared with the corresponding control groups, the GST activity in the AcT1 (8.69 IU/L) and AcT2 (7.67 IU/L) groups was significantly increased (*p* < 0.05), while that in the AcT3 group (7.50 IU/L) was reduced (*p* > 0.05) (Figure 5).

**Figure 5.**
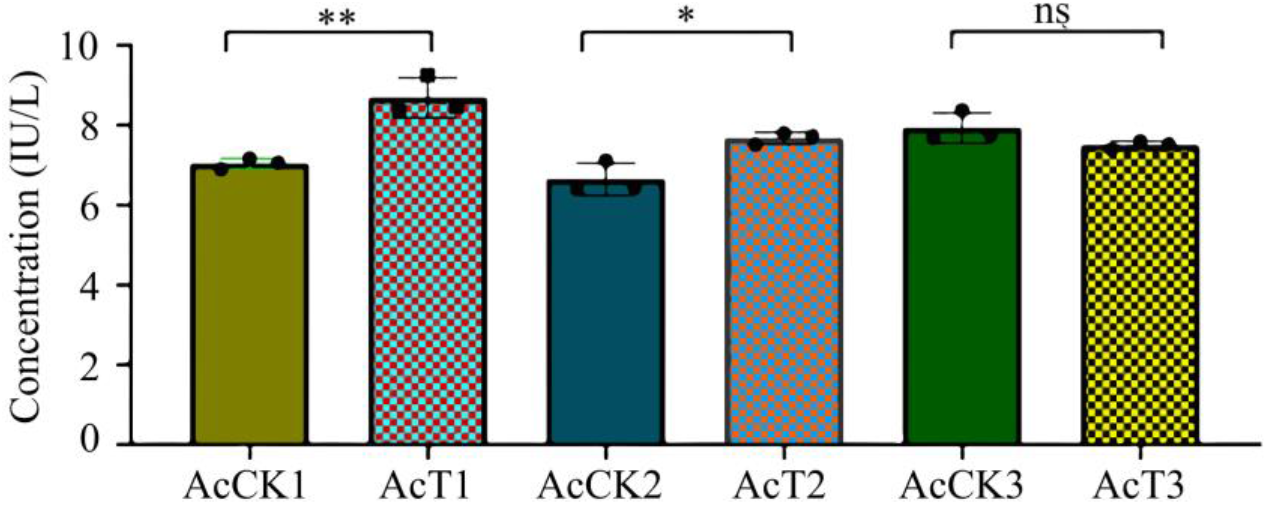
GST activity in *A. cerana* 4-, 5-, and 6-day-old larval guts infected by *A. apis*.

### 3.6. Effect of *A. apis* infection on PPO activity in *A. cerana* larval guts

Compared with the corresponding control groups, AcT1 PPO activity (80.08 U/g) was increased (*p* > 0.05), AcT2 PPO activity (99.27 U/g) was significantly increased (*p* < 0.05), and AcT3 PPO activity (8.74 U/g) was significantly decreased (*p* < 0.01), as shown in Figure 6.

**Figure 6.**
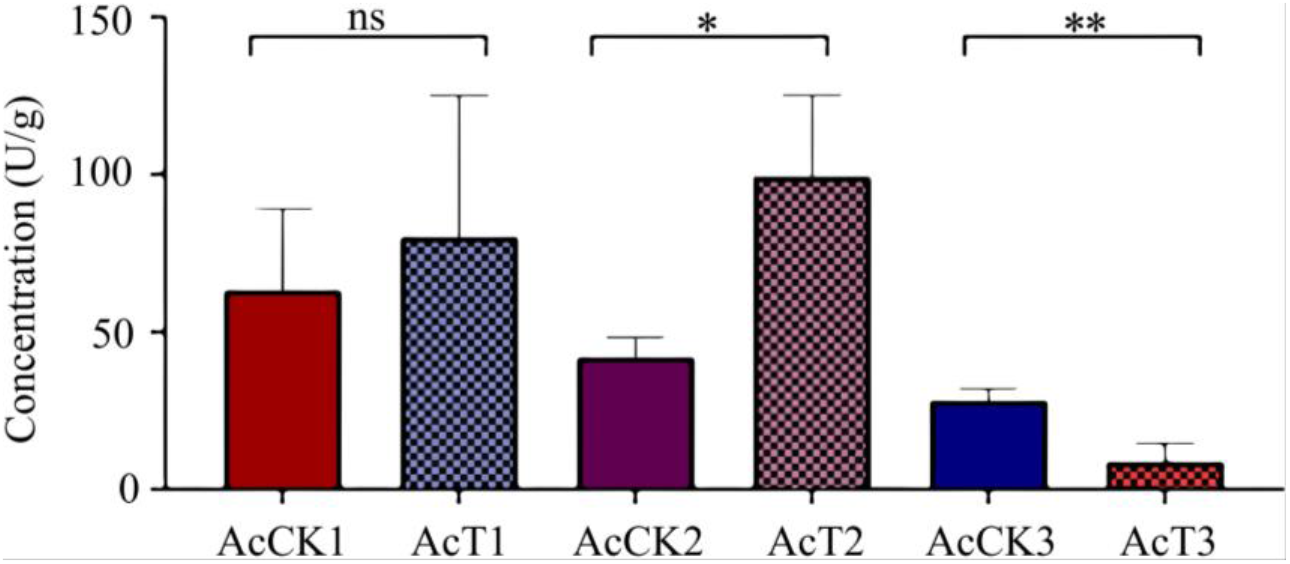
PPO activity in *A. cerana* 4-, 5-, and 6-day-old larval guts infected by *A. apis*.

## 4. Discussion

We have documented that *A. apis* spores were consumed by *A. mellifera* larvae via food sharing and germinated in the midgut lumen, with the disappearance of the diaphragm between the midgut and hindgut at the prepupal stage. The fungal spores and food debris swarmed into the hindgut lumen, where the mycelia rapidly grew in contact with O_2_, followed by penetration of the peritrophic membrane, gut wall, and body wall, eventually covering the entire larva, with a thick layer of white mycelia [4]. In the current work, the *A. apis* spores were purified and mixed with the diet to feed 3-day-old larvae. No apparent symptoms of chalkbrood disease were detected at 3–6 dpi. However, at 4 dpi, mycelia penetrated from the posterior end of the larva and then extended to the anterior end, eventually covering the entire larval body surface (Figure 1A). This course of chalkbrood disease was similar to that of *A. mellifera* larvae infected by *A. apis* [22], which was in line with the fact that *A. apis* was an exclusive fungal pathogen of bee larvae. Additionally, agarose gel electrophoresis showed that fragments of expected sizes (about 217 bp) could be amplified from *A. apis*-inoculated larval guts and *A. apis* spores, but not from the un-inoculated larval guts and sterile water (Figure 1B). This confirms the *A. apis* infection of *A. cerana* larvae after spore inoculation under lab conditions.

Here, we observed that the survival rate of *A. cerana* larvae after *A. apis* inoculation was 97.92%, 83.33%, 33.33% at 1~6 dpi, and sharply decreased to 4.16% at 4 dpi and 0 at 5 dpi, while that of un-inoculated larvae was always high at 1~8 dpi (95.37% on average) (Figure 2). This indicated that increased infection time of *A. apis* negatively influenced larval survival, following the pathogenesis mentioned above. To the best of our knowledge, this is the first experimental evidence of the survival rate of *A. cerana* larvae infected by *A. apis*.

In insects, protective and detoxifying enzymes are critical in maintaining normal physiological functions and biochemical metabolisms [23,10,24]. Among these, SOD and CAT can remove oxidative and toxic molecules such as O^2-^ and hydroxyl radical OH^-^ produced by exogenous compounds [11]. Li et al. [10] reported that both SOD and CAT activities were significantly reduced in the 3-day-old *A. m. ligustica* workers’ larvae at 96 h post inoculation with *A. apis* spores. In this work, we found that compared with the corresponding un-infected groups, the SOD and CAT activity in the 5- and 6-day-old *A. apis*-infected larval guts was significantly decreased (*p* < 0.05) (Figure 3, Figure 4), similar to the finding in the *A. m. ligustica* larvae infected by *A. apis* [10]. However, the SOD and CAT activities in the 4-day-old larval guts were reduced (*p* > 0.05), but there was no significant difference between *A. apis*-infected and un-infected groups (Figure 3, Figure 4), showing little influence of the two aforementioned antioxidant enzymes since there was only low-level spore germination and mycelial growth in the early stage *A. apis* infection. Together, these results demonstrated that *A. apis* infestation could negatively influence the SOD and CAT activities of honey bee larvae, which may be a strategy employed by *A. apis* during the long-term coevolution and interaction. After feeding the fourth and fifth-instar larvae of *Hyphantria cunea* with the leaves of *Bacillus thuringiensis* transgenic poplar, Ding et al. [25] detected that the activities of both SOD and CAT displayed an increase–decrease trend. Zhang et al. [24] inoculated *Nilaparvata lugens Stål* with *Metarhizium flavoviride* spores and found that SOD and CAT activities continuously increased as infection time prolonged. Furthermore, the SOD and CAT activities exhibited different trends in various insects responding to pathogen infections.

As a key component of the antioxidant enzyme system, GST is involved in the detoxification process of exogenous compounds in honey bee [13,14. Yan et al. [26] discovered that infection by highly pathogenic strains of entomo pathogenic nematode significantly altered the GST activity in the *Anoplophora glabripennis* larvae, presenting an overall increase–decrease–increase trend. Huang et al. [27] observed that GST activity was significantly higher in *Plutella xylostella* L. parasitized by *Diadegma semiclausum* than in the control group. In the present study, we found that GST activity in 4- and 5-day-old larval guts after *A. apis* infestation was significantly increased, as shown in Figure 5. Thus, *A. cerana* larvae were likely to enhance GST activity in response to oxidative stress caused by *A. apis* infestation. In insects, PPO is engaged in melanin formation, keratinization, and wound healing and also exerts a pivotal function in host immune defense [28,29]. The activity and content of PPO are often used as an indicator for evaluating insect immunity [29,24]. Li et al. [30] documented that PPO activity in infected *Apriona germari* larvae hemolymph increased to a maximum at 2.5 d post *Beauveria bassiana* infection and then decreased at 3.0 d. Wertheim et al. [31] reported that the expression level of the *PPO3* gene in *Drosophila* was significantly up-regulated at 48–72 h after the parasitic wasp challenge. In *Nilaparvata lugens Stål*,there was a significant elevation of PO content at 72 h post infection with *Metarhizium flavoviride* [24]. Here, we observed increased PPO activity in the 4-day-old larval gut, which was significantly elevated in the 5-day-old larval gut after *A. apis* infestation. The results showed that with the accumulation of spores and mycelia in the larval gut, the host reinforced the PPO activity to combat the *A. apis* infestation. Intriguingly, the PPO activity was significantly decreased in the gut tissue of 5-day-old larvae infected by *A. apis* (Figure 6). This indicated that at the late stage of infection, host PPO activity was suppressed due to the growing fungal stress. Taken together, these findings were suggestive of complex interactions between *A. cerana* larvae and *A. apis*.

## 5. Conclusions

Taken together, the *A. apis* spore inoculation of *A. cerana* larvae gave rise to chalkbrood disease, which decreased the host survival rate. The *A. apis* infestation affected the activities of SOD, CAT, GST, and PPO, suggestive of a host–adopted defending strategy mediated by antioxidant enzymes and complex host-pathogen interactions.

**Figure 7.**
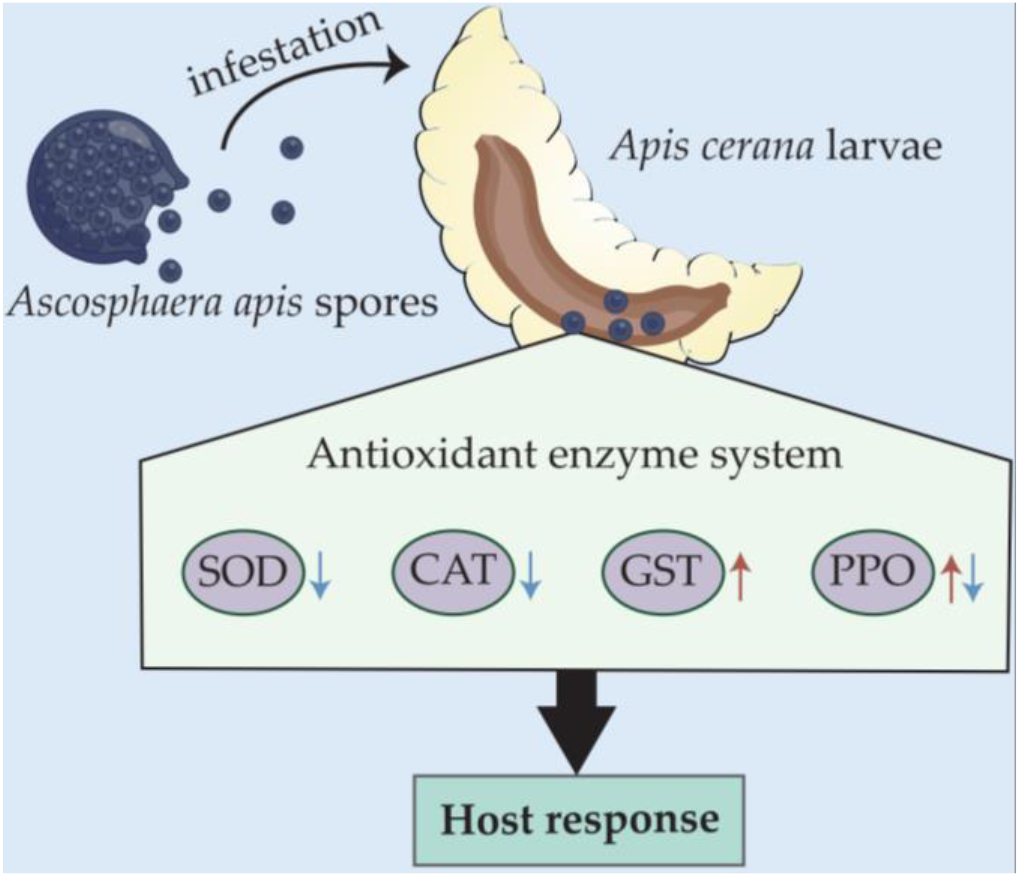
A hypothetical schematic diagram of the effect of *A. apis* infestation on the activities of four antioxidant enzymes in Asian honey bee larval guts.

## Author Contributions

R.G. and D.C. designed this research; K.Z., Z.F. contributed to the writing of the article; K.Z., Z.F., X. F., Z.W., S.W., S.G., X.G., H.Z., X.J., P.Z., Q.L. and M.C. conducted experiments and data analyses. R.G. and D.C. supervised the study and preparation of the manuscript.

## Acknowledgments

We thank all editors and reviewers for their constructive comments and recommendations. RG appreciates his adorable daughter for her love and goodness.

